# Lesions in a songbird vocal circuit increase variability in song syntax

**DOI:** 10.1101/2023.07.25.550508

**Authors:** Avani Koparkar, Timothy L. Warren, Jonathan D. Charlesworth, Sooyoon Shin, Michael S. Brainard, Lena Veit

## Abstract

Complex motor skills like speech and dance are composed of ordered sequences of simpler elements, but the neuronal basis for syntactic ordering of individual actions into sequences is poorly understood. Birdsong is a learned vocal behavior composed of syntactically ordered sequences of individual syllables. Activity in song premotor nucleus HVC (proper name) has been linked to the control of syllable sequencing, but sequencing may also be affected by its recurrent inputs. We here test the contribution of one of HVC’s inputs, mMAN (*medial magnocellular nucleus of the anterior nidopallium*), to the variable songs of adult male Bengalese finches (*Lonchura striata domestica*). The syntax of Bengalese song includes several patterns: 1) *chunks*, where syllables follow stereotypical order 2) *branch points*, where a given syllable can be followed by two or more different syllables in a probabilistic manner and 3) *repeat phrases*, where an individual syllable is repeated a variable number of times. We found that after bilateral lesions of mMAN, the acoustic structure of syllables remained largely intact, but sequencing became more variable for each of these patterns, seen by ‘breaks’ in previously stereotyped chunks, increased uncertainty at branch points and increased variability of repeat numbers. This increase in sequencing variability highlights the potential importance of regions projecting to HVC in the ordering of vocal elements. Previous studies on zebra finches found little effect of mMAN lesions on their relatively stereotyped adult song. In contrast, our results suggest that mMAN contributes to sequencing the variable songs of Bengalese finches and highlight the utility of species with more complex song syntax in investigating neuronal control of ordered motor sequences.

## Introduction

Complex behaviors are composed of sequences of simpler motor elements. For example, in human speech, sentences are formed by syllables that are flexibly sequenced according to syntactic rules (Berwick *et al*., 2011), which define the order of individual elements. Production of motor sequences is important not only for speech, but also for most other natural movements. Despite the ubiquity of syntactically organized sequential behaviors, little is known about how the brain produces flexible motor sequences (Lashley, 1951; Tanji, 2001; Aldridge and Berridge, 2003; Wiltschko *et al*., 2015; Veit *et al*., 2021).

Birdsong is a learned behavior that is well suited for the study of the neural control of motor sequences. Song is composed of discrete elements, called syllables, that are organized according to syntactic rules. The complexity of song syntax varies widely across different songbird species - from the simple and repetitive songs of owl finches (Wang *et al*., 2019) and zebra finches (Zann, 1996) to the variable songs of Bengalese finches (Honda and Okanoya, 1999) and canaries (Nottebohm, Stokes and Leonard, 1976; Cohen *et al*., 2020) up to the immense repertoire of nightingales (Hultsch *et al*., 2004; Costalunga *et al*., 2023). The neuronal mechanisms of birdsong production have been primarily studied in the zebra finch, partly by leveraging the structure of zebra finch song, in which syllables are produced in a relatively stereotyped and linear sequence. Consequently, we have a detailed understanding of the neuronal mechanisms for producing individual song syllables in this species, but we know little about how syllables are organized into the more complex and variable sequences that are characteristic of many other species (Ivanitskii and Marova, 2022).

Song production is controlled by neuronal activity in a set of brain nuclei called the song motor pathway, including premotor nucleus HVC (proper name). HVC neurons projecting to primary motor nucleus RA (*robust nucleus of the arcopallium*) encode syllable identity and have been shown to play a role in controlling syllable timing (Simpson and Vicario, 1990; Vu, Mazurek and Kuo, 1994; Hahnloser, Kozhevnikov and Fee, 2002; Long and Fee, 2008; Aronov *et al*., 2011; Ölveczky *et al*., 2011; Lynch *et al*., 2016; Picardo *et al*., 2016; Zhang *et al*., 2017). HVC activity in Bengalese finches (Sakata and Brainard, 2008; Fujimoto, Hasegawa and Watanabe, 2011) and canaries (Cohen *et al*., 2020) has also been shown to encode syllable identity, and can additionally influence sequencing and reflect the sequential context in which syllables are produced (Fujimoto, Hasegawa and Watanabe, 2011; Zhang *et al*., 2017; Cohen *et al*., 2020). The nuclei of the motor pathway are essential for song production (Nottebohm, Stokes and Leonard, 1976; Scharff and Nottebohm, 1991), but the extent to which these sequence-selective activity patterns in HVC reflect internal neural dynamics or are shaped by recurrent inputs is poorly understood.

Although HVC is often considered to be at the top of a hierarchy in the song control circuit, it is part of a network of brain nuclei involved in song production and receives multiple inputs that might also contribute to premotor activity (McCasland, 1987; Hosino and Okanoya, 2000; Jin, 2009; Hamaguchi, Tanaka and Mooney, 2016; Vyssotski *et al*., 2016). One prominent source of input to HVC is mMAN (*medial magnocellular nucleus of the anterior nidopallium*) which is part of a recurrent loop that bilaterally connects the two hemispheres through RA and bilateral connections from thalamic nucleus DMP (*dorsomedial nucleus of the posterior thalamus,* Figure 1E) and could thus be ideally positioned to convey sequence-related activity across hemispheres (Vates, Vicario and Nottebohm, 1997; Schmidt, Ashmore and Vu, 2004; Williams, Nast and Coleman, 2012). A parallel recurrent pathway, which projects to RA through nucleus lMAN (*lateral magnocellular nucleus of the anterior nidopallium*, Figure 1E) has been shown to contribute variability to syllable pitch, a parameter that is controlled by RA premotor activity (Kao, Doupe and Brainard, 2005; Sober, Wohlgemuth and Brainard, 2008; Miller, Cheung and Brainard, 2017). lMAN-guided variability can serve as a form of motor exploration which is essential for learning (Sober and Brainard, 2012; Dhawale, Smith and Ölveczky, 2017), regulating social context-dependent changes in pitch variability (Kao and Brainard, 2006; Hampton, Sakata and Brainard, 2009), and biasing pitch in the direction of improved motor performance during learning (Andalman and Fee, 2009; Warren *et al*., 2012; Tian and Brainard, 2017). This raises the question of whether mMAN, as the output of a parallel recurrent pathway that projects onto HVC, might have a similar role in contributing to premotor activity in its projection target, with the potential to influence syllable identity or sequencing variability (Kubikova, Turner and Jarvis, 2007; Seki and Okanoya, 2008; Ali *et al*., 2013). Previous studies in zebra finches showed that mMAN lesions in juveniles disrupt normal song learning, while lesions in adults affected song initiation, but not subsequent production of song in this species (Foster and Bottjer, 2001; Horita, Wada and Jarvis, 2008; Ali *et al*., 2013).

**Figure 1:**
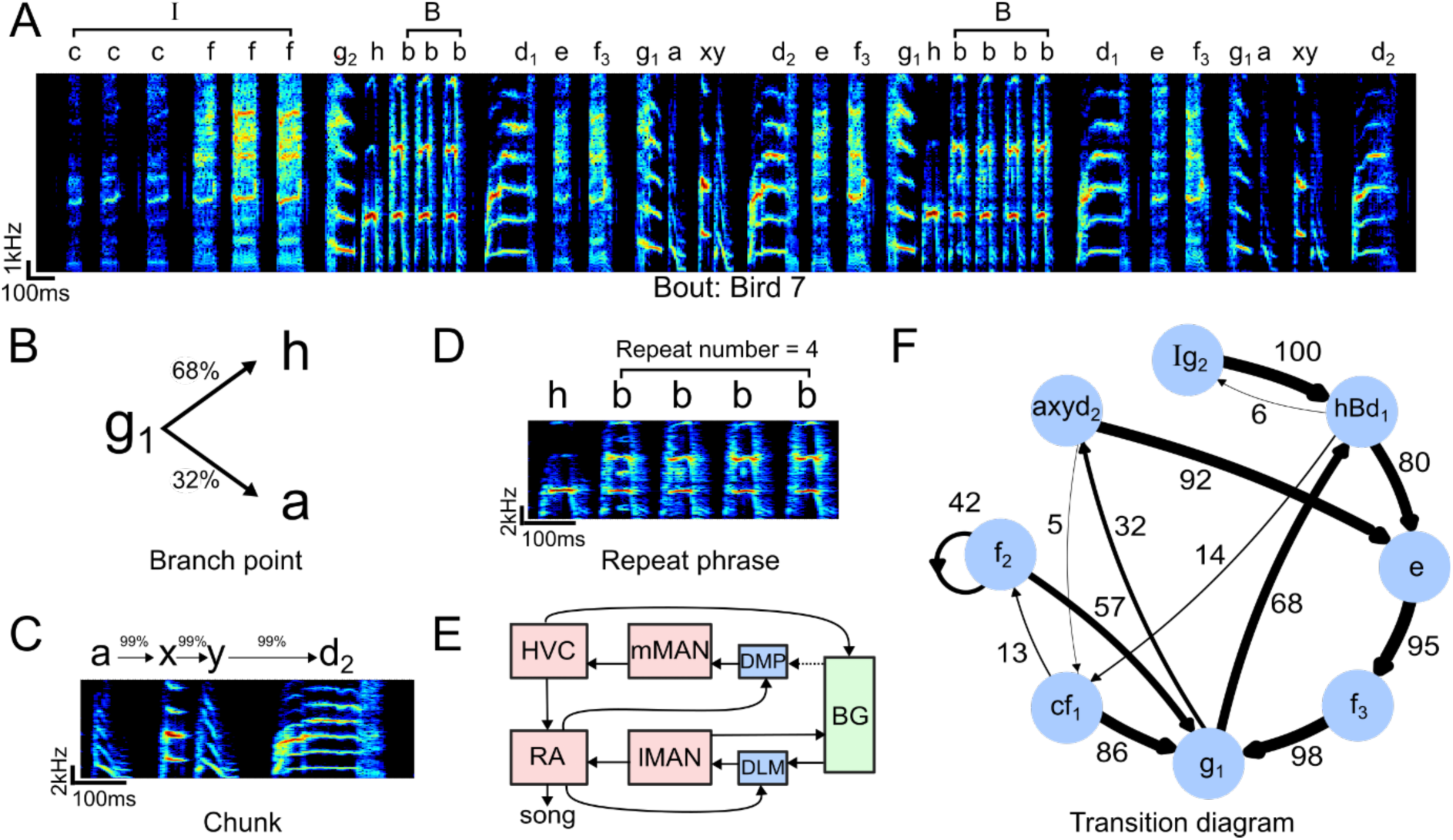
Structure of Bengalese finch song (A) Example spectrogram (bird 7) depicting an entire song bout with introductory state *‘I’*, repeat phrase ‘*B*’ and individual syllables. (B) Example transition diagram depicting a branchpoint with variable sequencing. Numbers above the arrows denote transition probabilities in percent. (C) Example spectrogram of a chunk. Chunks are defined as highly stereotyped sequences of syllables which only have a single in- and output branch and are condensed into one state in the transition diagram (see Methods). (D) Example spectrogram of a repeat phrase, summarized by capital letter ‘*B*’ in the transition diagram. The repeating syllable (here, syllable *‘b’*) repeats a variable number of times across different instances of the repeat phrase. (E) Schematic showing recurrent pathways projecting onto motor pathway nuclei through lMAN and mMAN. Red: pallial nuclei, blue: thalamic nuclei, green: basal ganglia. Dotted line indicates suspected connection by Kubikova et al. 2007. Abbrev: mMAN: *medial magnocellular nucleus of the anterior nidopallium*, DMP: *dorsomedial nucleus of the posterior thalamus*, BG: *basal ganglia*, DLM: *medial portion of the dorsolateral thalamus*, lMAN: *lateral magnocellular nucleus of the anterior nidopallium*, RA: *robust nucleus of the arcopallium*. (F) Example of a transition diagram. Nodes denote chunk or syllable labels, numbers denote transition probabilities (in percent, % symbol omitted for clarity), *d1*/*d2*, *g1*/*g2* denote different states of syllables *d* and *g* respectively based on different sequential contexts, capital letters denote repeat phrases. Edges at each node may not sum to 100% because branches smaller than 5% are omitted for clarity.

We tested the hypothesis that mMAN contributes to syllable sequencing by studying the effect of bilateral mMAN lesions on song production in adult Bengalese finches. We found that lesions had little effect on the acoustic structure of individual syllables but led to an increase in the variability of multiple aspects of syllable sequencing. These results highlight the potential importance of recurrent inputs such as mMAN in shaping the syntactical structure of adult birdsongs.

## Methods

### Subjects and sound recordings

Experiments were carried out on seven adult male Bengalese finches (*Lonchura striata domestica*) obtained from the Brainard lab’s breeding colony at University of California, San Francisco (median age 640 days post-hatch, range 149-1049 at start of experiment). Birds were raised with their parents and then housed in same-sex group cages. For the experiments, birds were placed in individual sound-attenuating boxes (Acoustic Systems, Austin, TX) and maintained on a 14:10 hour light: dark period. Song was recorded using an omnidirectional microphone above the cage using custom LabView software for continuous monitoring of song output (Tumer and Brainard, 2007). All procedures were performed in accordance with animal care protocols approved by the University of California, San Francisco Institutional Animal Care and Use Committee (IACUC).

### mMAN lesions

After several days of sound recording, birds were anesthetized using ketamine and midazolam with isoflurane. mMAN was localized using stereotaxic coordinates and was destroyed bilaterally using current injections of 100uA current for 80s at four sites per hemisphere. After conclusion of the study, birds were deeply anesthetized and perfused with paraformaldehyde. 40um sections were cut on a microtome and processed with either Nissl stain (cresyl violet) or calcitonin gene-related peptide (CGRP) (Hampton, Sakata and Brainard, 2009). mMAN lesion size was estimated by experienced observers and estimated to be either complete (bird 1, birds 3-6) or greater than 75% (bird 2 and 7) for all birds used in this analysis (Figure 1-figure supplement 1).

### Sequence analysis

Bengalese finch songs were recorded for several days before and after lesions. Singing activity resumed on average 3.8 days post lesions (range 2-6 days, Figure 1-figure supplement 1), and we analyzed songs starting on average 5.3 days post lesions (range 3-11 days) up to 7.4 days post lesions (range 5-12 days). We analyzed at least two days pre and post lesions, and subsampled the larger of the two datasets pooled over all days such that the pre and post datasets were equal in size for each bird (average of 299 song bouts, range 102-601, for all analyses). All our main analyses were carried out on this dataset. To analyze specifically the persistence of effects after the lesion, Figure 1-figure supplement 2 includes four additional data points (14, 19, 33 and 33 days post lesion for birds 2, 3, 6 and 7 respectively) after additional behavioral manipulation, from which birds typically recover (Warren *et al*., 2012).

**Figure 2:**
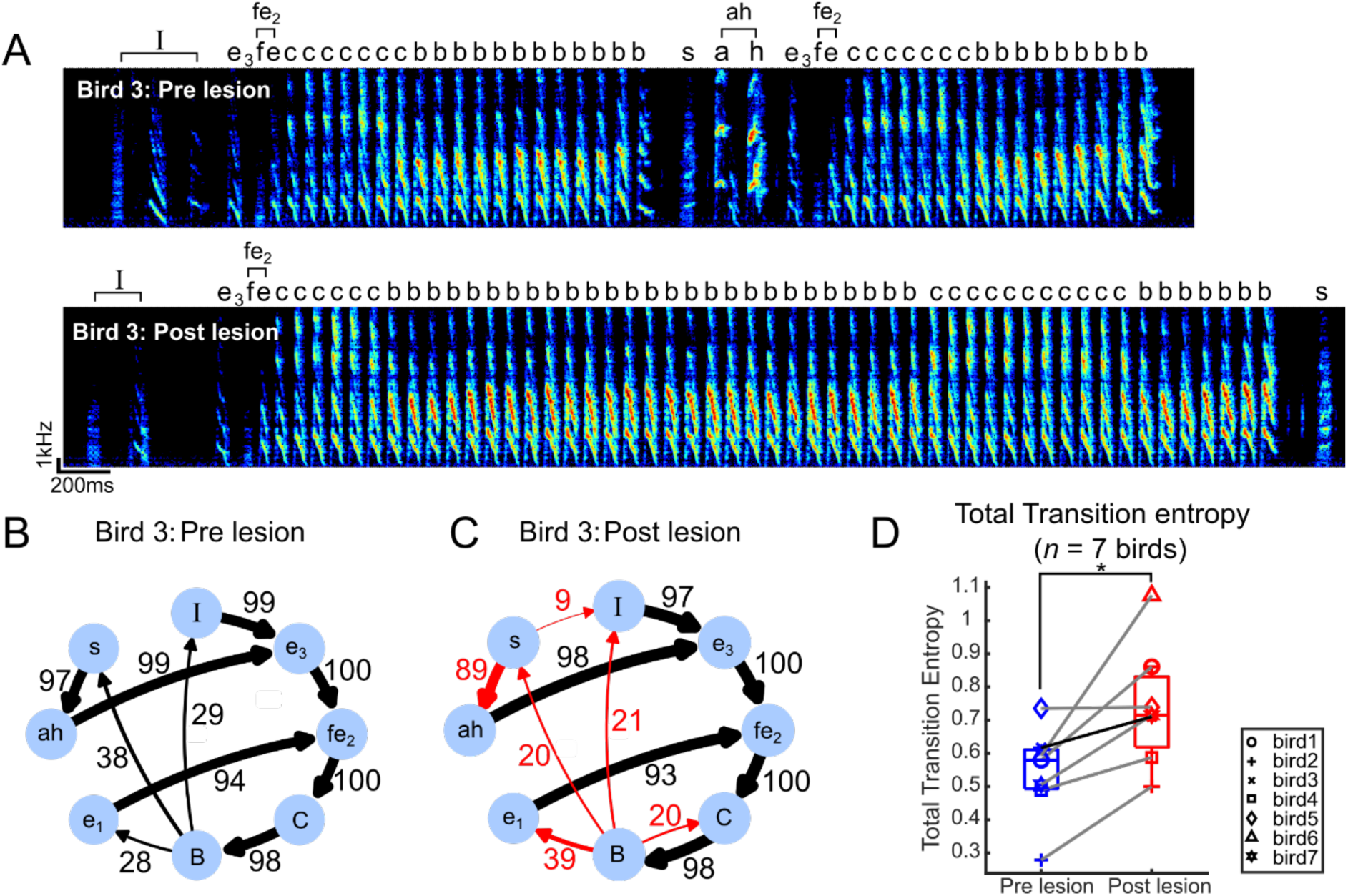
Transition entropy increased after bilateral mMAN lesions (A) Example spectrogram (bird 3) pre lesion (above) and post lesion (below). *‘I’* denotes introductory state, and ‘*fe2’* and ‘*ah’* denote chunks, shown as single nodes in the transition diagram. (B) Pre lesion transition diagram, as in 1F. Note that *‘fe2CB’* is also a chunk before the lesions but is shown as three separate nodes in order to align with the post lesion diagram in C. (C) Post lesion transition diagram. Arrows in red mark example nodes with relatively high increase in transition entropy, including the introduction of new branches after mMAN lesions. (D) Total transition entropy for 7 birds before and after mMAN lesions (* p<0.05, n=7, Wilcoxon signed-rank test). Example bird from A-C is shown as darker line.

#### Annotation

We annotated the entire syllable sequence in an automated way. We first used a subset of the data to generate a training set for classification. Separate classifiers were trained for each bird. In the training data, syllable onsets and offsets were determined by amplitude thresholding. We then performed dimensionality-reduction and unsupervised clustering of the spectrograms of detected syllables using UMAP (McInnes *et al*., 2018; Sainburg, Thielk and Gentner, 2020), in order to determine the number of different syllables in an objective way (mean 13, range 11-14). On average, we used 181 files (range 70-505) for creating this training dataset, which was then used to train a deep neural network (TweetyNet) for the annotation of birdsong (Cohen *et al*., 2022). TweetyNet was used to segment and annotate all songs, followed by semi-automated hand-checking using custom-written software in MATLAB R2021b to ensure quality annotation. In rare cases, where there was some ambiguity in the assignment of syllable identity during hand-checking, we additionally took into account the sequential context, for example during stereotyped chunks, to assign the most likely label. Thus, any errors in assignment in such cases would have tended to reduce rather than accentuate the magnitude of the lesions’ effects on reported sequencing changes.

#### Simplification of the syllable sequence

We defined song bouts as continuous sequences of syllables separated by at least 2 s of silence. We then simplified the syllable sequence by merging introductory notes into a single introductory state, i.e., a single node ‘*I*’ in the transition diagram (Figure 1A,F). Introductory notes were defined as up to three syllable types occurring at the start of bouts which were quieter, and more variable in timing and structure than other syllable types (Rajan and Doupe, 2013; Veit *et al*., 2021). Similarly, syllables which were repeated a variable number of at least two times were reduced to a single state (e.g. ‘*bbbbbbbbb*’ to ‘*B*’, Figure 1A,F) in the syllable sequence, and the number of repetitions within these repeat phrases was analyzed separately (Hampton, Sakata and Brainard, 2009; Zhang *et al*., 2017; Jaffe and Brainard, 2020). We only analyzed repeat phrases with variable numbers of syllable repetitions in this way, therefore, we did not consider cases in which a syllable was repeated an exact number of times on each occurrence (occurring with repeat numbers of 2 or 3 in our dataset, see Figure 3A) a variable repeat phrase.

**Figure 3:**
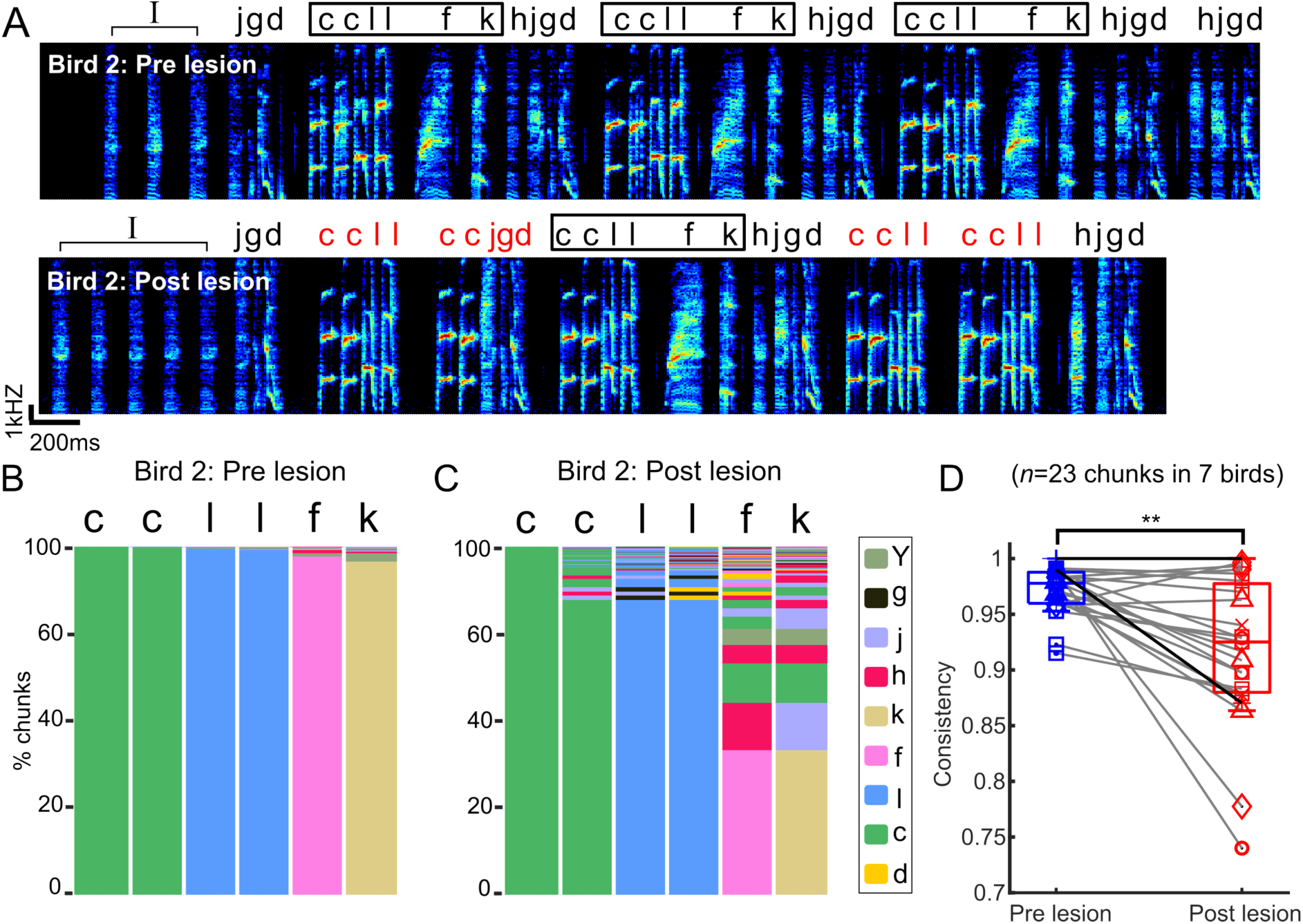
Chunks became more variable after bilateral mMAN lesions. (A) Example spectrogram (bird 2) before and after bilateral mMAN lesions. Atypical chunk sequences are highlighted in red. (B,C) Transitions following the first ‘*c*’ of the *‘ccllfk’* chunk from (A) before and after mMAN lesions. Different column colors represent different syllables. (D) Chunk consistency before and after bilateral mMAN lesions (** p<0.01, n=23, Wilcoxon signed rank test). Example bird is shown as darker line.

#### Separating different syllable states based on sequential context

Transition probabilities between syllables in Bengalese finch song can depend not only on the identity of the syllable, but also on the identity of at least one preceding syllable (Jin and Kozhevnikov, 2011; Katahira *et al*., 2011; Morita *et al*., 2021). In order to consider this sequential context-dependency of transition probabilities, we followed the procedure of Katahira et al. 2011 to classify spectrally similar syllables (i.e., with the same labels) into different ‘states’ based on the identity of the preceding syllable. For example, in Figure 1A, the syllable *‘g’* is preceded by *‘I’* or *‘f’* and is followed by *‘h’* or *‘a’*. If *‘g’* is preceded by *‘I’* it is always followed by *‘h’* and if it is preceded by *’f’*, it can be followed by *‘h’* or *‘a’*. We would therefore like to introduce two separate states *g_1_* and *g_2_* in the transition diagram representing this sequential context. The input for this analysis was the syllable sequence with introductory and repeat phrases already condensed into single states. We compared the distribution of transition probabilities from these syllables by themselves (e.g., syllable *‘g’*) to the distribution of transition probabilities from these syllables considering the context of one preceding syllable (e.g., syllable *‘f-g’*) using the chi-square goodness of fit test. We split the syllable into a new state (e.g. *‘g_1_’*) if *p* < *0.01/n* where *n* is the number of comparisons made (Bonferroni correction for multiple comparisons) (Jin, 2009; Katahira *et al*., 2013). After completing this for all syllables of a given type (e.g., *‘g_1_’*, *‘g_2_’*, *‘g_3_’*), we compared the different output states to each other using chi-square goodness of fit test and merged them back together if *p > 0.01/n* (mean 15, range 6-21 additional states/bird).

#### Determining chunks and branchpoints

To determine chunks (Suge and Okanoya, 2010), we used the modified syllable sequence after chi-square analysis. We calculated a syllable-to-syllable transition diagram, while eliminating any syllables occurring with less than a threshold frequency of 0.9% and omitting branches with a probability of less than 5%. From the resultant graph, we merged all non-overlapping linear sequences (which consisted of nodes with only a single input and output branch) into chunks. The transition probabilities within resultant chunks were typically higher than 90% (range 87-100%). Values smaller than 100% for non-branching paths result from the omission of one or several small branches of less than 5%. We then re-calculate the transition probabilities between chunked nodes. We defined branchpoints as the set of single syllables that are retained after this processing and the end syllables of the newly defined chunks. To simplify diagrams for visualization, we eliminated any syllables which occurred less than once per bout and branches with transition probabilities of less than 5%.

#### Transition entropy

Transition entropy is a measure of uncertainty of sequence at a given syllable (Sakata and Brainard, 2006). With *c* different outputs from the given syllable *‘a’* and *P(i)* the probability of the *i^th^* outcome, we calculate the entropy *H_a_,* as:

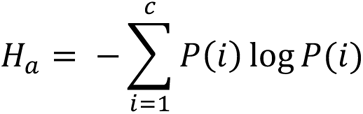

We call this value ‘transition entropy per branchpoint’ in Figure 3A.

To determine the overall variability of song before and after mMAN lesions, we calculated total transition entropy *TE*, over all syllables *‘b’* as:

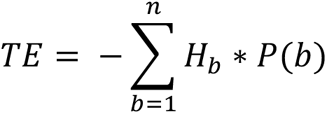

where *H_b_* is the transition entropy at *‘b’* and *P(b)* is the frequency of syllable *‘b’* (Chatfield and Lemon, 1970; Katahira *et al*., 2013).

#### History dependence

History dependence is a previously established metric that measures the extent to which the most common transition at a given syllable is influenced by the transition at the last occurrence of this syllable (Warren *et al*., 2012). It has been used to characterize instances of apparent sequence variability, where seemingly variable transitions are always strictly alternating. For example, if the possible transitions from syllable *‘a’* are *‘ab’* or *‘ac’* but these strictly alternated (*‘ab…ac…ab…ac’* and so on) then the seemingly variable branchpoint *‘a’* is perfectly predictable based on its history (Warren *et al*., 2012). Such apparent variability should be largely eliminated in our sequence analysis by the introduction of context dependent states (i.e., in this example, the *‘a’* would be re-labelled as *‘a_1_’* or *‘a_2_’* depending on the context in which it occurs) and identification of chunks. However, if higher-order dependencies in the song determine the order of chunks, we might still expect some variable transitions to be governed by history dependence. If *‘ab*’ is the most frequently occurring transition from *‘a’*, and *‘ac’* is the collection of all other transitions from *‘a’*, we define history dependence D of *‘a’* as:

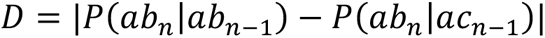

where 𝑃(𝑎𝑏*_n_*|𝑎𝑏*_n_*_-1_) is the conditional probability of *‘ab’* transition given that *‘ab’* transition occurred at the previous instance of *‘a’* and 𝑃(𝑎𝑏*_n_*|𝑎*c_n_*_-1_) is the conditional probability of *‘ab’* transition given that *‘ac’* transition occurred at the previous instance of *‘a’*.

#### Chunk consistency

As defined above, a chunk is defined by a single dominant sequence, but may have a small amount of variability across different instances. To quantify the stereotypy of chunks, we used a measure based on sequence consistency previously defined for relatively stereotyped zebra finch songs (Scharff and Nottebohm, 1991). Across all instances of a given chunk, we identified the syllable sequence that occurred most often as the ‘dominant sequence’. We then defined ‘*n_dominant*’ as the number of instances of the dominant sequence, and ‘*n_other*’ as the number of instances of other sequence variants for the chunk.

We quantified chunk consistency *C* as the proportion of total instances that were the dominant sequence:

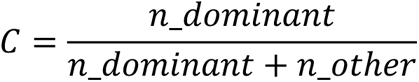

To compare a chunk before and after mMAN lesions, the dominant sequence for the pre lesion chunk was used as a reference, regardless of whether the same sequence qualified as a chunk post lesion. To quantify chunk consistency post lesion, the most dominant sequence post lesion was used (even if that was not the same as the most dominant sequence pre lesion).

#### Repeat number variability

To study the influence of mMAN on repeat phrases, we examined the distribution of repeat numbers before and after lesions. We quantified the variability of these distributions as their coefficient of variation (*CV = standard deviation/mean*).

### Analysis of syllable acoustic structure

#### Average spectrogram

For visual inspection of preservation of syllable types before and after mMAN lesions, we calculated average spectrograms for 200 instances of each syllable type for 7 birds (Figure 2-figure supplement 1). Individual spectrograms of each syllable type were rescaled to have matching time axes and then overlaid.

#### Syllable similarity analysis

We used Sound Analysis Pro (SAP) software (Tchernichovski *et al*., 2000) to quantify the similarity in syllable structure for all syllable types per bird (Figure 2-figure supplement 2). We used the settings ‘Symmetric’ and ‘Time Courses’ and compared twenty instances of each syllable type.

We first calculated SAP similarity of all syllables to syllables of the same type from two separate control recordings before mMAN lesions. We termed this as ‘Self Similarity’. We next calculated SAP similarity of the same syllable types before and after mMAN lesions. We termed this as ‘Pre vs Post’ similarity. Lastly, we calculated SAP similarity of all syllable types compared to all other syllable types per bird. This was used as a measure of dissimilarity of structure across syllable types. We termed this as ‘Cross Similarity’.

#### Analysis of fundamental frequency

We used the method previously described by Hampton, Sakata and Brainard, 2009 to calculate fundamental frequency and coefficient of variation (CV) of fundamental frequency for select syllables for each bird (Figure 2-figure supplement 3) (Hampton, Sakata and Brainard, 2009). Briefly, each syllable was visually inspected to have a segment with clearly defined fundamental frequency. We then found the peak of the autocorrelation function (using parabolic interpolation) of the sound waveform in this segment.

#### Analysis of acoustic features

To further compare syllable phonology before and after the mMAN lesions, we used the SoundSig python package (Elie and Theunissen, 2016) to calculate a set of acoustic features for all syllables for 7 birds (Figure 2-figure supplement 4). The features used were Entropy of spectral envelope (entS), Temporal centroid for the temporal envelope (meanT), first, second and third formants (F1, F2, F3).

## Results

### Structure of Bengalese finch song

Bengalese finch songs are composed of a fixed number of syllable types, which are arranged into variable sequences following syntactical rules. In this study, we sought to first characterize the different features of Bengalese finch song, before investigating their change following mMAN lesions. For each bird, we first annotated all songs with labels for each of the syllable types, and then defined a transition diagram for analysis according to the following procedure (see Methods):

1) We identified introductory notes (Rajan and Doupe, 2013; Veit *et al*., 2021) and grouped them together into a single introductory state *‘I’* in the transition diagram. This reduced the overall transition entropy of song for 7 birds from 0.95 to 0.85 (Figure 1A).
2) Repeat phrases, in which the same syllable is repeated a variable number of times, were also reduced to a single state in the transition diagram (indicated by capital letters) and the distribution of repeat numbers was analyzed separately (Hampton, Sakata and Brainard, 2009; Zhang *et al*., 2017; Jaffe and Brainard, 2020) (Figure 1D).
3) In Bengalese finch song a single spectrally defined and labelled syllable can occur in different fixed sequences (for example, syllable *‘b’* in the sequences *a-b-c* and *d-b-e*) and therefore reflects distinct states in the song sequence (Wohlgemuth, Sober and Brainard, 2010; Katahira *et al*., 2011). We followed the procedure of Katahira et al. to determine if any initially identified syllable types corresponded to more than one state based on whether the preceding syllable carried information on transition probabilities (see Methods, Katahira *et al*., 2013). After identifying and splitting these ‘hidden states’ (for example *b_1_*, *b_2_*) for each bird, overall transition entropy was further reduced from 0.85 to 0.54 (Figure 1F).
4) In the new syllable sequence, we identified ‘chunks’ (Suge and Okanoya, 2010), sub-sequences of high-probability transitions. We defined these as paths of continuous, non-branching sequences in the transition diagram, after omitting branches that occurred with probability < 5% (Figure 1C). Lastly, we identified branch points in the transition diagram as the set of single syllables and endpoints of chunks, i.e., all states where the syllable sequence proceeds in a probabilistic manner (Figure 1B,F).

### Transition entropy increased after bilateral mMAN lesions

After bilateral mMAN lesions, all syllable types were preserved (Figure 2-figure supplement 1-4), but the sequencing of syllables became more variable. In the example in Figure 2B,C, we show the transition diagrams of one example bird before and after mMAN lesions. We observed a change in transition probabilities at existing branch points, as well as the appearance of novel branches. We quantified changes in sequencing variability for all syllables using transition entropy (Sakata and Brainard, 2006; Katahira *et al*., 2013). Total transition entropy increased significantly after mMAN lesions, indicating that song syntax overall became more variable (Figure 2D). These effects appeared as soon as the bird resumed singing after the lesions and did not recover over the time for which the birds were observed (Figure 1-figure supplement 2).

Figure 2-figure supplement 5A shows the corresponding changes at all individual branch points. To investigate the source of differences in the magnitude of changes across branch points, we considered the history-dependence of each of these branchpoints. History-dependence captures apparent variability governed by long-range dependencies in the sequence and history-dependent transitions previously have been found to be difficult to modify in a sequence modification training protocol (Warren *et al*., 2012). They might therefore be less affected by lesions as well. Alternatively, we might expect lesion effects to be stronger for these transitions, if mMAN contributes specifically to long-range dependencies. Consistent with the first possibility, we observed that there was a nonsignificant trend toward larger changes after mMAN lesion for transitions with low history dependence. (Figure 2-figure supplement 5B).

### Chunks became more variable after bilateral mMAN lesions

Bengalese finch songs contain short, relatively stereotyped sequences of syllables, here called ‘chunks’ (Okanoya and Yamaguchi, 1997; Suge and Okanoya, 2010; Isola, Vochin and Sakata, 2020; Veit *et al*., 2021). We speculated that transitions within chunks might be differently affected by mMAN lesions, if for example the relatively fixed sequences within chunks are determined within premotor song nucleus HVC and inputs to HVC are only relevant at variable branch points. We found that the relatively fixed sequences within chunks were altered after mMAN lesions. An example bird (Figure 3A-C) exhibited one prominent chunk which was occasionally observed to ‘break’ after lesions, i.e., branching within the chunk increased to a degree where it would no longer be considered a chunk after the lesions. The branching increased most noticeably in the *‘l-f’* transition (Figure 3C), which is characterized by a longer gap than other transitions within the chunk (Figure 3A). We therefore wondered whether sequencing changes were related to gap durations and found that gap durations within chunks were not significantly correlated with the increase in transition entropy at the corresponding transitions (Figure 3-figure supplement 1A). Overall, we found that changes within chunks were approximately the same magnitude as changes at branch points (Figure 3-figure supplement 1B).

We quantified changes in within-chunk transitions using sequence consistency (Scharff and Nottebohm, 1991), a measure previously used to describe the relatively consistent sequence of zebra finch song. Sequence consistency measures how consistently the most probable sequence is followed (see Methods). Sequence consistency within chunks significantly decreased across 23 chunks from all birds, indicating that this aspect of song structure was affected by mMAN lesions (Figure 3D).

### Repeat numbers became more variable after mMAN lesions

We next tested how mMAN lesions affected repeat phrases. Repeat phrases might be governed by separate neural mechanisms than branch points (Fujimoto, Hasegawa and Watanabe, 2011; Jin and Kozhevnikov, 2011; Wittenbach *et al*., 2015; Zhang *et al*., 2017). In our dataset of 7 birds, only 5 birds had songs which contained repeat phrases.

In the example bird in Figure 4A,B, the average repeat number for the indicated syllable increased after mMAN lesions. Across birds, the mean repeat number did not change consistently, although individual birds could exhibit quite dramatic effects on repeat number, (Figure 4B,C; p>0.05, n=6, Wilcoxon signed rank test). The distributions of repeat numbers became wider and the coefficient of variation for repeat numbers increased significantly after mMAN lesions (Figure 4D; * p<0.05, n=6, Wilcoxon signed rank test).

**Figure 4:**
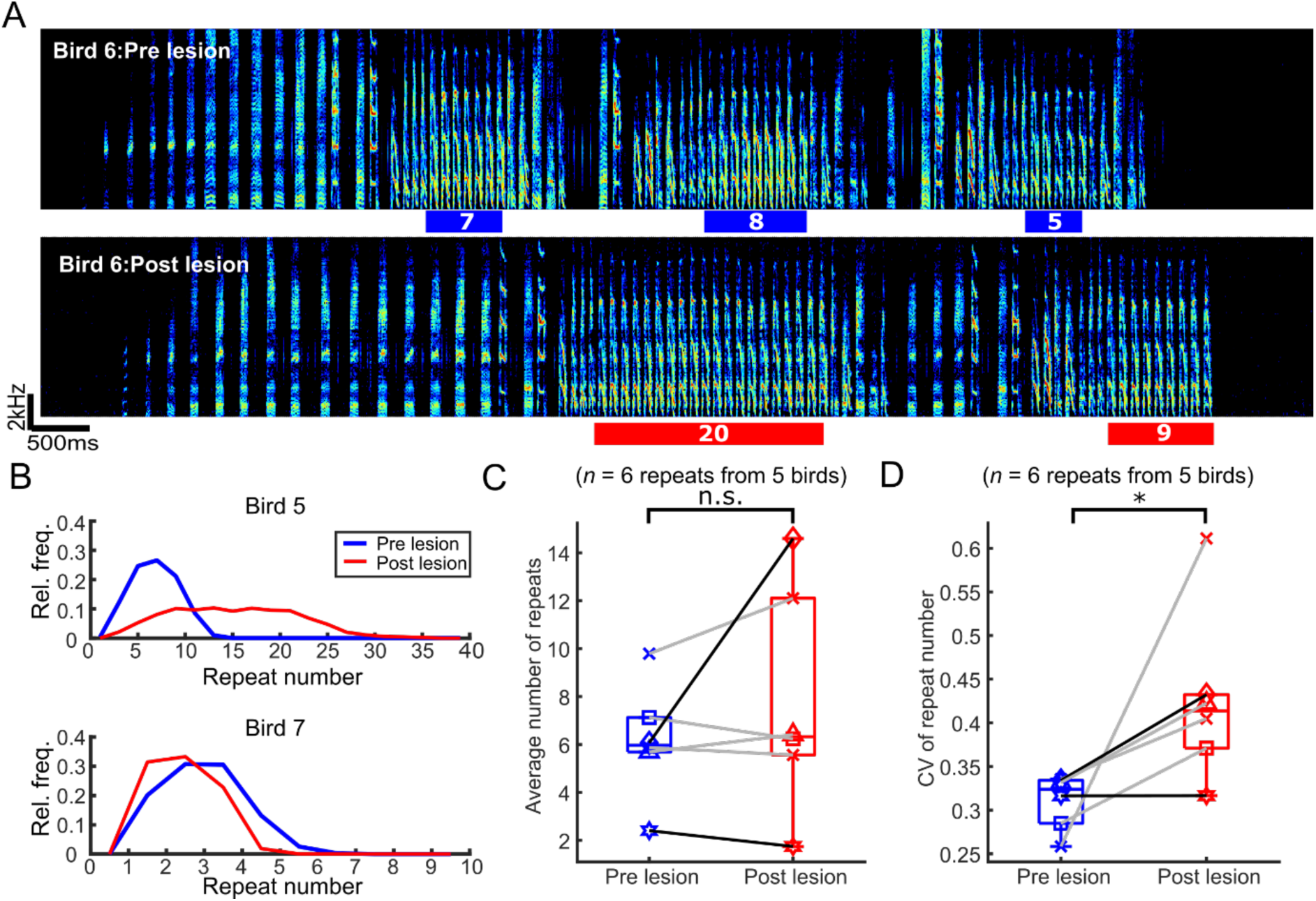
The number of syllables per repeat phrase (repeat number) became more variable after bilateral mMAN lesions. (A) Example spectrogram (bird 6) highlighting one repeat syllable before (blue) and after (red) mMAN lesions. (B) Repeat numbers for two additional example birds before (blue) and after (red) mMAN lesions. (C) Average repeat numbers before and after mMAN lesions for all repeat phrases. (D) Coefficient of variation for distribution of repeat numbers before and after mMAN lesions for all repeat phrases. Example birds from B are shown as darker lines.

## Discussion

Bird songs are vocal sequences composed of individual syllables which appear in sequential order dictated by syntactic rules, but the neuronal control of song syntax is largely unknown. Previous studies in zebra finches showed that mMAN lesions in adult birds had minimal effects on song production (Foster and Bottjer, 2001; Horita, Wada and Jarvis, 2008; Ali *et al*., 2013). We speculated that the modest effect of lesions in adult zebra finches might be due to the relatively stereotyped nature of zebra finch song. Consistent with this possibility, our results demonstrate that lesions of pallial nucleus mMAN affect the more complex and variable song syntax in Bengalese finches. We showed that while mMAN lesions did not abolish the production of song itself or grossly disrupt the acoustic structure of individual syllables, they increased the variability of syllable sequencing; the transition probabilities between syllables became less predictable (increased transition entropy), new branches were introduced into previously stereotyped chunks, and the number of repetitions of repeated syllables (repeat number) became more variable. These results indicate that mMAN is involved in controlling aspects of song syntax in a songbird species that exhibits complex syllable sequencing and suggest that mMAN might play a similar or prominent role in additional species with even greater song complexity, such as canaries or nightingales.

Song in passerine birds relies on a network of recurrently connected nuclei, with different pathways for song production and learning. The song motor pathway consists of primary motor nucleus RA, which projects directly to vocal motor nuclei in the brainstem, and is involved in the moment-by-moment control of the structure of individual syllables (Vu, Mazurek and Kuo, 1994; Sober, Wohlgemuth and Brainard, 2008; Miller, Cheung and Brainard, 2017). Neurons in premotor song nucleus HVC that project to RA have been shown to encode syllable identity and contribute to syllable timing (Simpson and Vicario, 1990; Vu, Mazurek and Kuo, 1994; Hahnloser, Kozhevnikov and Fee, 2002; Long and Fee, 2008; Aronov *et al*., 2011; Ölveczky *et al*., 2011; Zhang *et al*., 2017). Consistent with a possible role of HVC in also controlling variable syllable sequencing, cooling HVC influences syntax in Bengalese finches (Zhang *et al*., 2017). However, lesions of one input to HVC – the nucleus Nif (*nucleus interfacialis of the anterior nidopallium*) – can also influence syllable sequencing in the Bengalese finch by making the sequence more stereotyped (Hosino and Okanoya, 2000; Cardin, Raksin and Schmidt, 2005; Otchy *et al*., 2015) raising the question of how much of the control of syntax relies on the internal dynamics of HVC, versus activity relayed to HVC by its inputs (Okanoya and Yamaguchi, 1997).

In this study, we were especially interested in the possibility that mMAN’s inputs to HVC might play an analogous role in modulating syllable sequencing to the role of lMAN’s inputs to RA in modulating syllable acoustic structure. In adult birds, lMAN’s inputs to RA have been shown to contribute variability to the pitch of individual syllables (Kao, Doupe and Brainard, 2005; Kao and Brainard, 2006; Stepanek and Doupe, 2010), regulate changes to pitch variability between different social contexts (Kao and Brainard, 2006; Hampton, Sakata and Brainard, 2009), and bias pitch in the direction of improved performance during learning experiments (Andalman and Fee, 2009; Warren *et al*., 2011; Tian and Brainard, 2017). The variability contributed by lMAN may facilitate motor exploration which forms the basis of pitch adaptations during learning (Andalman and Fee, 2009; Sober and Brainard, 2012; Dhawale, Smith and Ölveczky, 2017). These effects are mediated by top-down influences of lMAN on RA via a recurrent loop that passes through the songbird basal ganglia (Kao, Doupe and Brainard, 2005; Andalman and Fee, 2009; Kojima *et al*., 2018; Tian, Warren and Brainard, 2022), and are consistent with a role of RA in controlling aspects of syllable acoustic structure. In contrast to these effects on syllable structure, lesions of lMAN in adults have not been found to appreciably influence syllable sequencing (Hampton, Sakata and Brainard, 2009). However, because mMAN forms the output of a parallel recurrent loop that passes through the basal ganglia and projects to HVC, we hypothesized that analogous modulation of syllable sequencing during production and learning of song might be mediated by mMAN. Our finding that lesions of mMAN alter multiple aspects of syllable sequencing are consistent with such an influence on song syntax.

The finding that mMAN can influence song syntax raises the question of whether and how this pathway modulates syllable sequencing in the service of motor exploration, contextual modulation, and learning, in a fashion that is analogous to lMAN’s modulation of syllable acoustic structure. That mMAN lesions consistently altered sequence variability indicates a possible role in regulating variability for motor exploration. However, whereas lMAN lesions consistently *decrease* the variability of acoustic structure, we found that mMAN lesions generally *increase* the variability of syllable sequencing, suggesting that top-down contributions to the control of syntax may be different or more distributed than for a syllable feature such as pitch. One possibility is that several inputs to HVC could work synergistically to modulate the syntactical structure of Bengalese finch songs. Consistent with this view, a prior study in Bengalese finches found that lesions of Nif decreased sequence variability (Hosino and Okanoya, 2000), suggesting that Nif and mMAN could be playing opposing roles in regulating variability. Our findings also raise the question of whether the medial anterior forebrain pathway including mMAN, plays a similar role in learning to change syllable sequences during development (Foster and Bottjer, 2001; Lipkind *et al*., 2013) or adulthood (Warren *et al*., 2012; Veit *et al*., 2021). In our study, we only recorded song sequencing of male Bengalese finches singing in isolation. Social context, such as female-directed song, can also change song sequencing (Hampton, Sakata and Brainard, 2009; Chen, Matheson and Sakata, 2016). It would be interesting to test whether mMAN plays a role in the social context-modulated changes in sequencing (Jarvis *et al*., 1998), similar to how lMAN contributes to social context-modulated changes in syllable structure (Sakata, Hampton and Brainard, 2008). Further studies with careful manipulations of the basal ganglia-mMAN-HVC circuit will be required to elucidate the mechanism with which this circuit affects sequencing in Bengalese finch song and contributes to these behaviors.

Bengalese finch syllable sequences contain structure at multiple levels, mainly the organization of individual stereotyped chunks, resembling the stereotyped motifs of zebra finch song, and the variable branch points between chunks, repeat phrases, and other individual syllables. One possibility to reconcile findings that both dynamics within HVC and its recurrent inputs affect sequencing could be that the sequence within chunks is encoded predominantly in the synaptic connections of HVC neurons (Jin, 2009), while the branching transitions found in Bengalese finch song are influenced by input from other nuclei (Hosino and Okanoya, 2000). We here show that mMAN lesions affect both transition entropy at previously variable branch points and transitions within previously stereotyped chunks, indicating that such a simple split of function between mMAN and HVC is not the case.

Besides these, other parts of the song circuit have been shown to affect syllable sequencing. The basal ganglia are involved in selection and sequencing of motor elements in mammals, and manipulations of songbird basal ganglia Area X have previously been shown to affect syllable repetitions in zebra finches and Bengalese finches (Kobayashi, Uno and Okanoya, 2001; Kubikova *et al*., 2014; Tanaka *et al*., 2016; Xiao *et al*., 2021). These effects might be partly mediated via connections through mMAN to HVC (Kubikova, Turner and Jarvis, 2007). Additionally, the control of syllable sequencing must include bilateral coordination of the motor pathways in the two hemispheres (Schmidt, Ashmore and Vu, 2004; Wang *et al*., 2008). As the two hemispheres in birds are not connected by a corpus callosum, the only pathways mediating interhemispheric synchronization of the two HVCs are by bilateral connections through thalamic nuclei. One such pathway involves a projection from thalamic nucleus Uva to HVC (Coleman and Vu, 2005; Hamaguchi, Tanaka and Mooney, 2016; Moll *et al*., 2023), which is required for song production and contributes to syllable initiation and timing. Another is through thalamic nucleus DMP, which projects bilaterally through mMAN to HVC (Schmidt, Ashmore and Vu, 2004; Seki and Okanoya, 2008). mMAN might therefore be well situated to coordinate syllable sequencing in the two hemispheres. Future studies will need to determine whether the effects we observed are the results of removing neuronal processing within mMAN itself, or whether mMAN is conveying signals to HVC from other brain areas, including the contralateral hemisphere, which contribute to the control of song syntax.

## Acknowledgements

We thank Jacqueline Göbl, Lioba Fortkord, Yarden Cohen, Tim Gardner, Felix Moll for helpful discussions and comments on earlier versions of this manuscript. We thank David Nicholson and Peter Pilz for help with technical questions. Michael Brainard was supported by the Howard Hughes Medical Institute. Lena Veit was supported as a Howard Hughes Medical Institute Fellow of the Life Sciences Research Foundation and by a Daimler Benz postdoctoral fellowship. Avani Koparkar was supported by the Studienstiftung des deutschen Volkes and International Max Planck Research School for Mechanisms of Mental Function and Dysfunction (IMPRS-MMFD). Jonathan Charlesworth and Timothy Warren were supported by the National Science Foundation (NSF) Graduate Research Fellowship. We acknowledge support from the Open Access Publication Fund of the University of Tübingen.

## Author contributions

T.L.W., J.C., S.S. and L.V. gathered the data. A.K. and L.V. analyzed the data, A.K., L.V. and

M.S.B. wrote the paper.

## Supplementary Figs

**Figure 1 - figure supplement 1:**
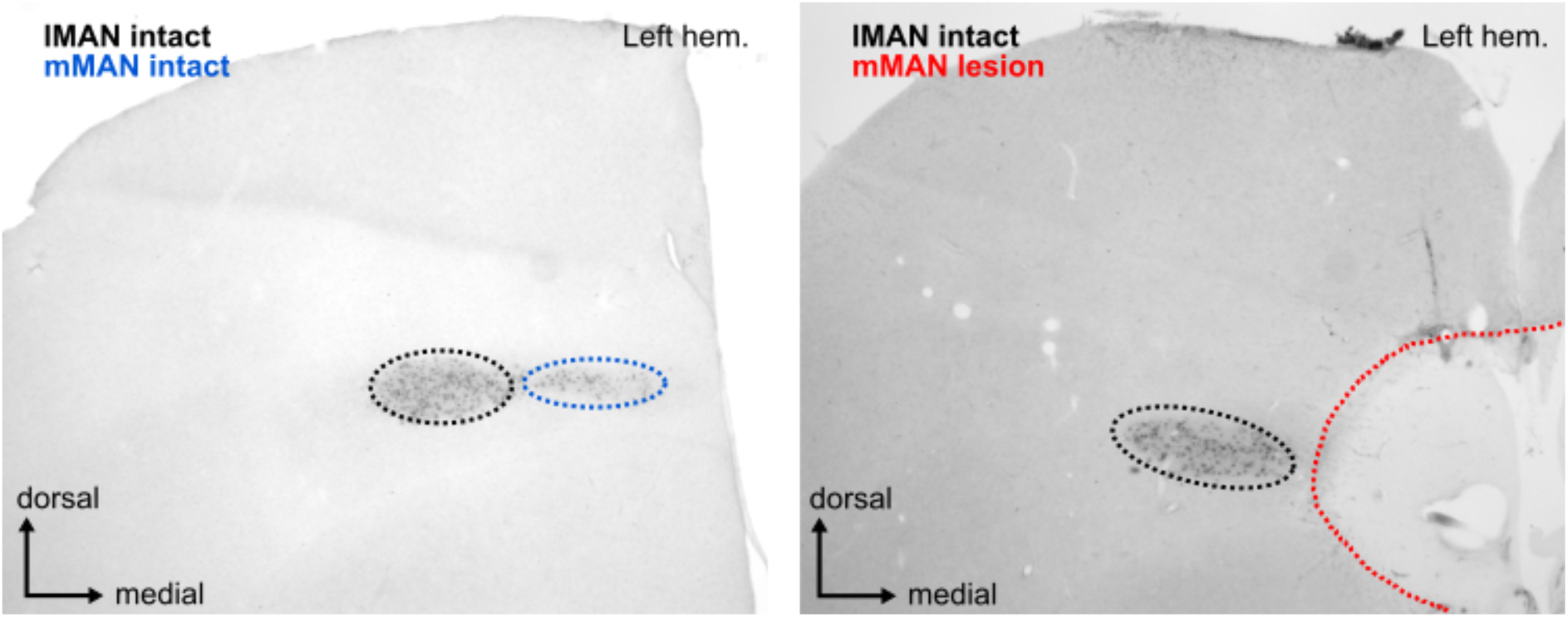
Image of calcitonin gene-related peptide (CGRP)-stained frontal section (left) control and (right) bird 5. CGRP labels cells in both lMAN (seen in black to the left of the lesion) and mMAN (blue, intact; red, completely destroyed).

**Figure 1 - figure supplement 2:**
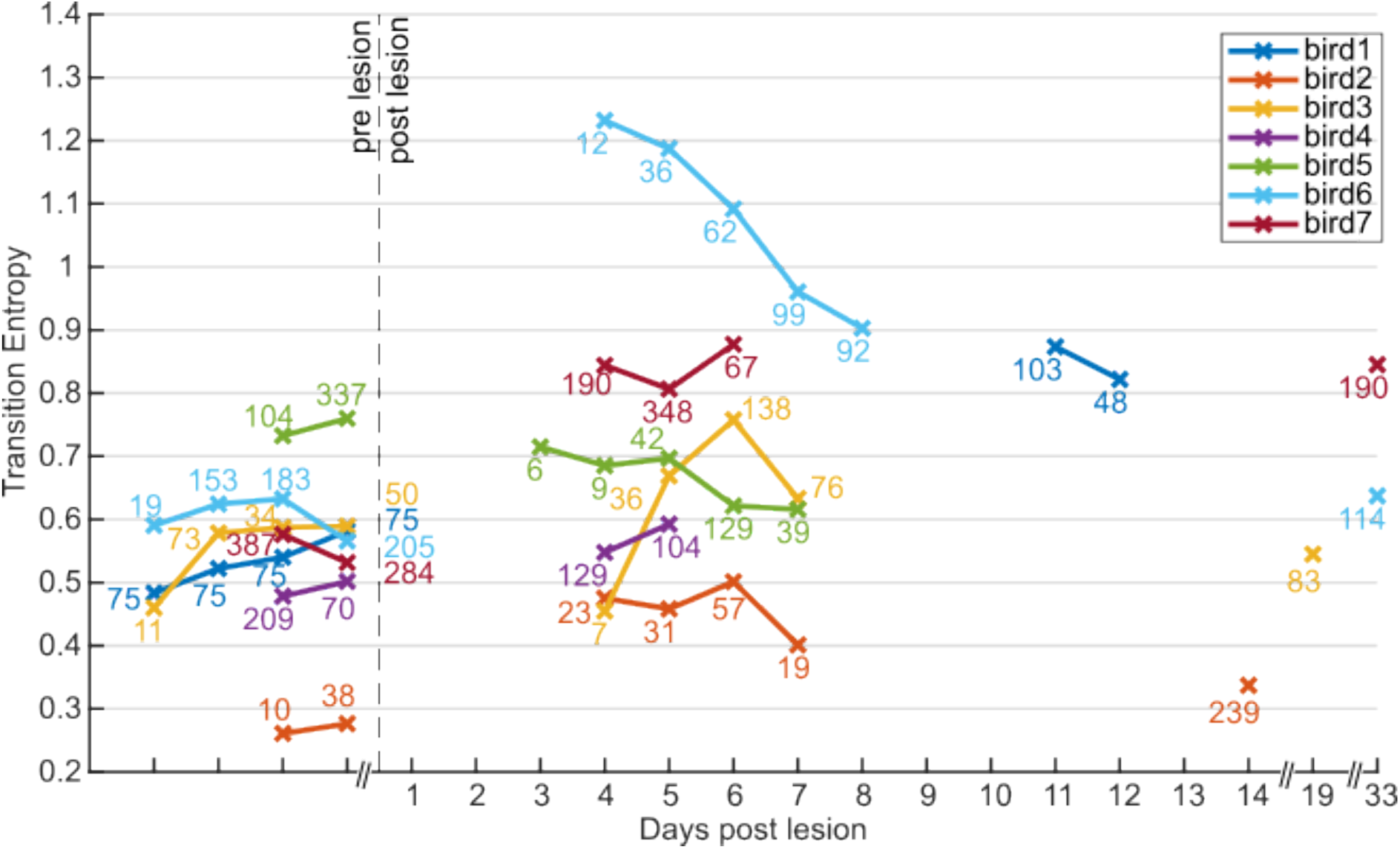
Transition entropy after mMAN lesions remains elevated over several days. Transition entropy did not change in a consistent way after recovery from the lesion for the followed time period. Numbers near the datapoints indicate number of song bouts recorded and analyzed for that day. While the persistence of the effects we observed is longer than transient effects such as those following Nif lesion in zebra finches (∼ 2 days by Otchy *et al*., 2015) we cannot rule out either recovery or further deterioration following lesions on much longer time scales, such as those reported by Kubikova et al., 2007 (X lesions, 6 months) (Kubikova, Turner and Jarvis, 2007). Additional non-connected datapoints at 14, 19, 33 and 33 days post lesion were gathered after additional behavior manipulation and were added here to analyze specifically the persistence of effects after the lesion. For three of these birds, transition entropy remained elevated above the baseline values for 14, 33 and 33 days, respectively.

**Figure 2 - figure supplement 1:**
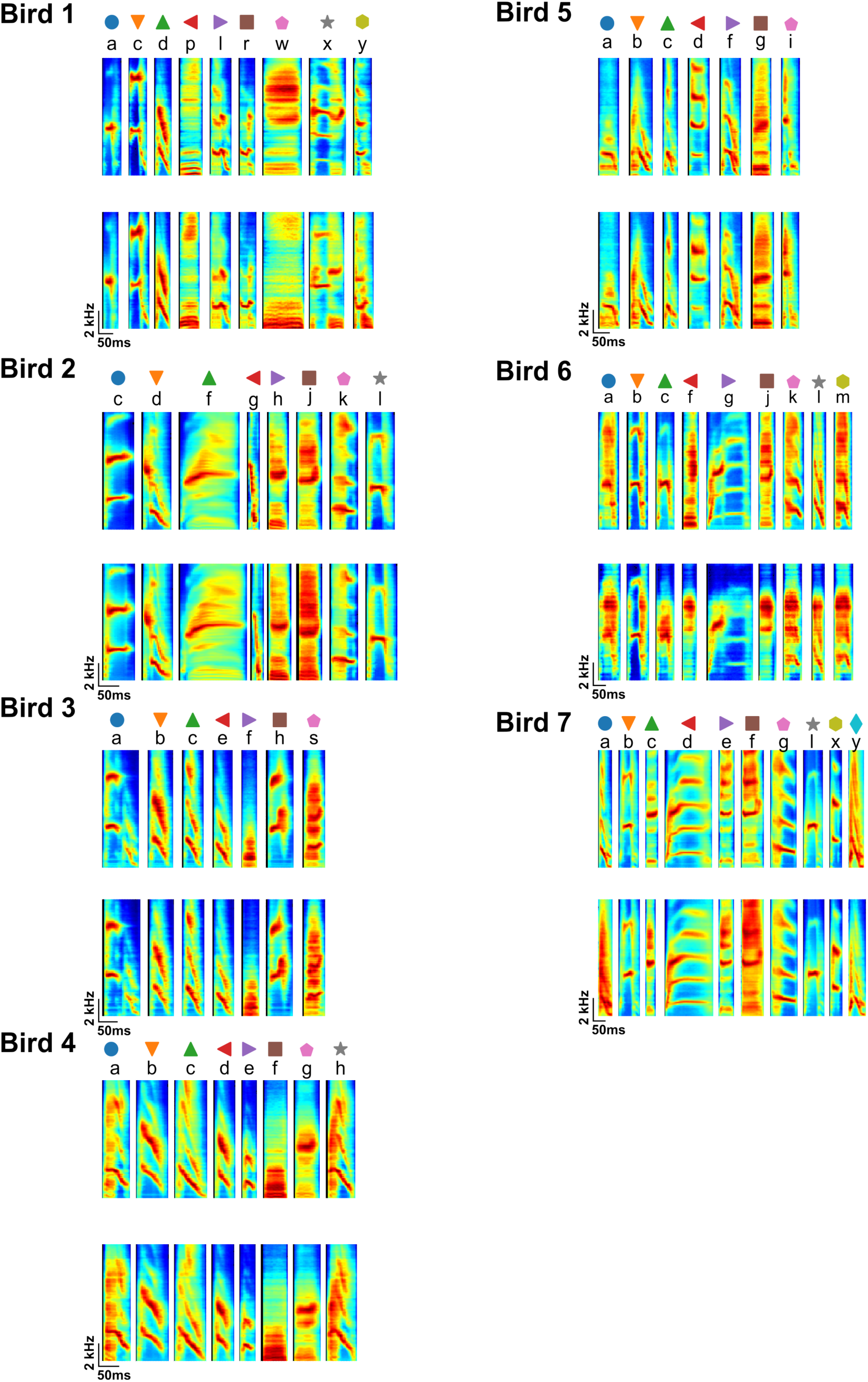
Average spectrograms of 200 instances of all syllable types for all birds before and after mMAN lesions. Letters over the spectrograms indicate syllable labels and symbols over the spectrograms correspond to the symbols used in Fig. 2 – figure supplement 4.

**Figure 2 - figure supplement 2:**
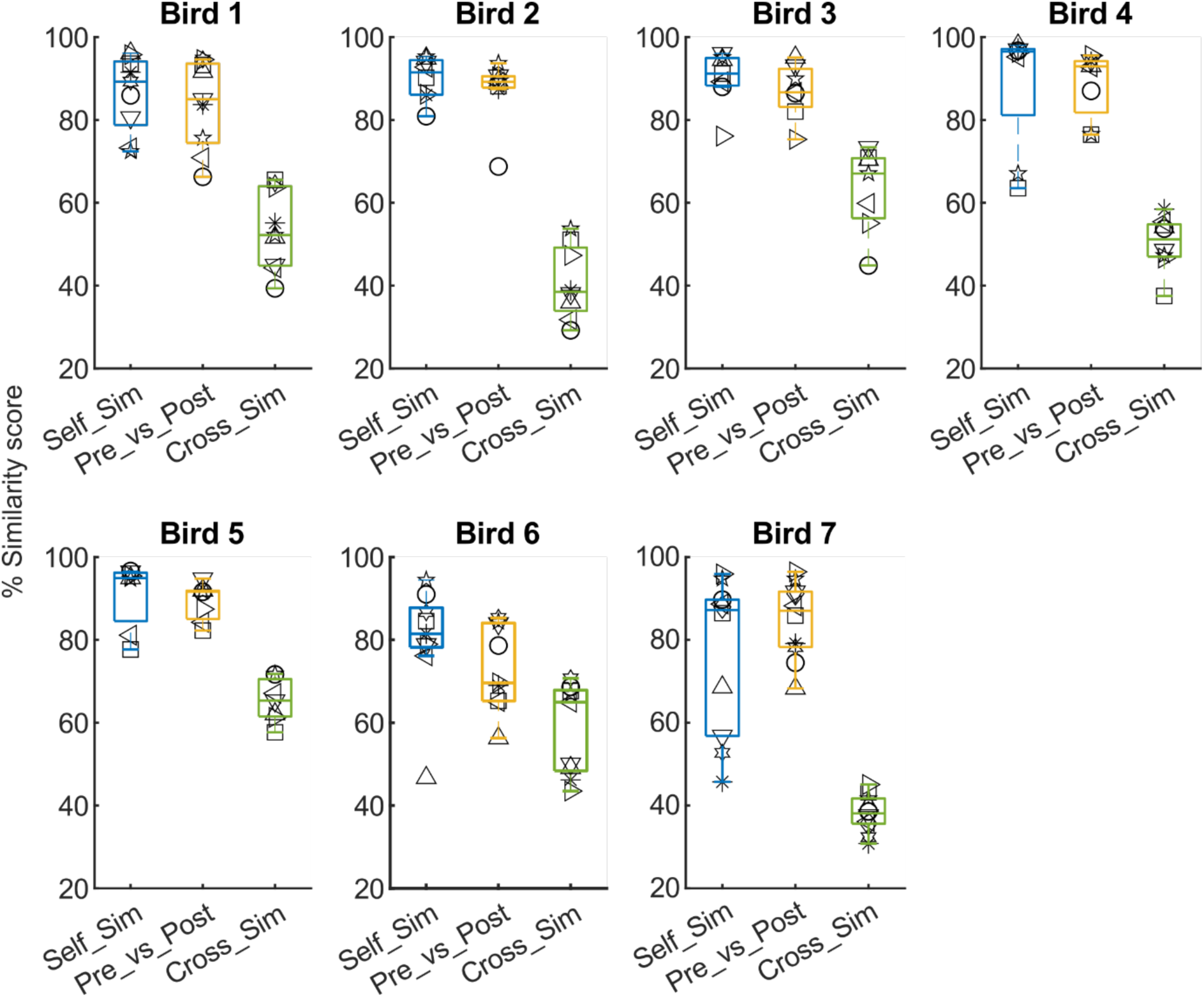
Syllable similarity calculated using Sound Analysis Pro (SAP). ‘Self Similarity’ = Similarity comparison of syllables before mMAN lesions to syllables of the same type, taken from two separate control recordings before the lesions, ‘Pre vs Post’ = Similarity comparison of the same syllable types before and after mMAN lesions, ‘Cross Similarity’ = Similarity comparison of each syllable type to other syllable types. For Birds 1-2 and 4-7, ‘Self Similarity’ was not significantly different from ‘Pre vs Post’ Similarity (p>0.05, Wilcoxon sign rank test), while for Bird 3, there was a significant difference (p = 0.03, Wilcoxon sign rank test). For all birds ‘Pre vs Post’ was significantly different from ‘Cross Similarity’ (p<0.05, Wilcoxon sign rank test). On average, ‘Pre vs Post’ was 4.8 % less than ‘Self Similarity’ (range 0.2%-14%) while ‘Cross Similarity’ was 40% less than ‘Self Similarity’ (range 20.2%-56.3%). These measures confirm the qualitative impression from Figure 2- figure supplement 1 that for most birds and syllables there were no greater changes to syllable structure following lesions than was present across control recordings, and that pre-post similarity remained higher than cross-similarity, i.e. syllables remained clearly identifiable.

**Figure 2 - figure supplement 3:**
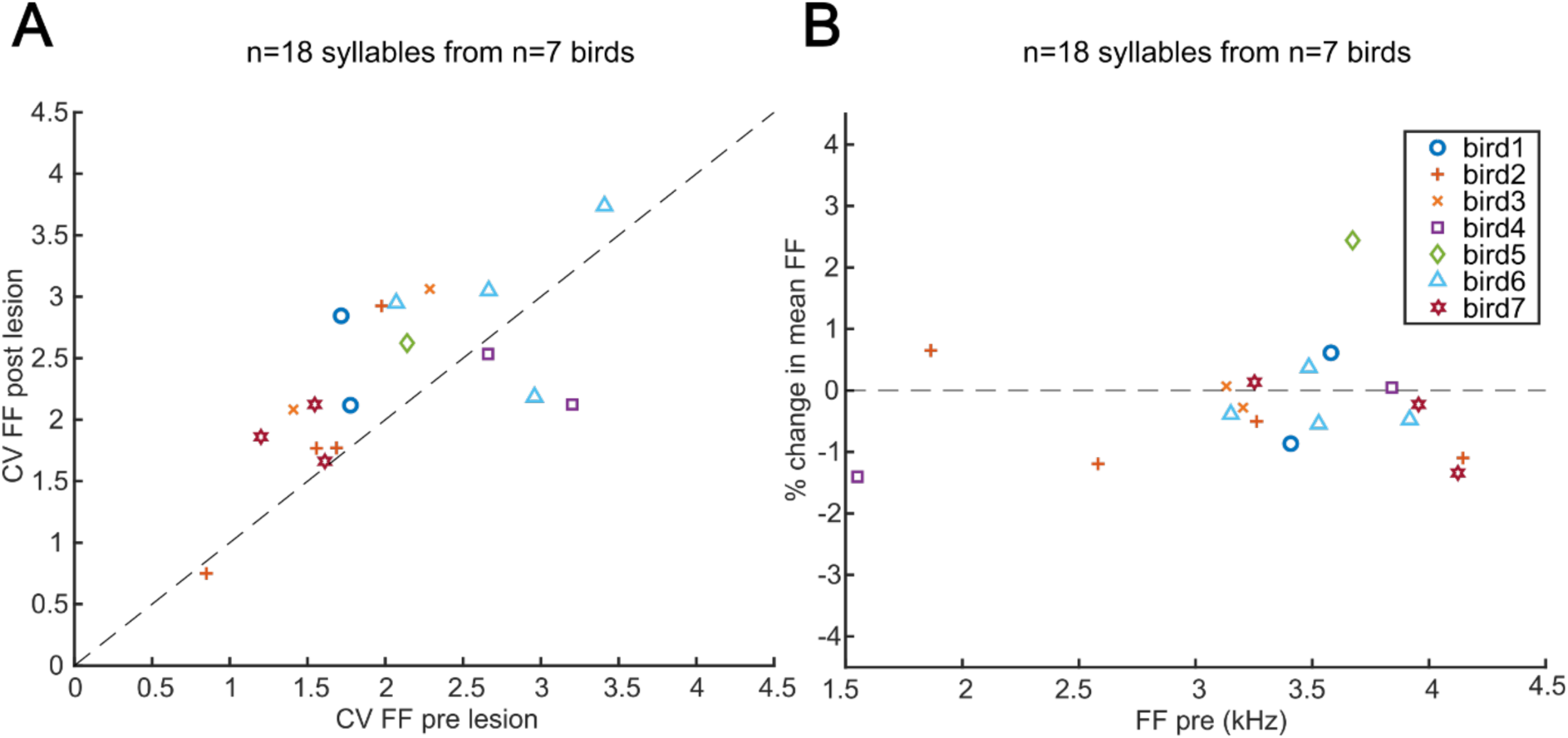
(A) CV of fundamental frequency (FF) of select syllables before and after mMAN lesions. In the Bengalese finch and zebra finch, lesions of lMAN, which sits immediately lateral to mMAN, cause a consistent reduction in the coefficient of variation (CV) of fundamental frequency across repeated renditions of a given syllable (Sakata, Hampton, Brainard 2008, Andalman, Fee 2009, Warren et al. 2011). We therefore supposed that unintended damage to lMAN or its projections to RA might have resulted in a reduction in the CV of syllables following mMAN lesions. Instead we saw a modest increase in the CV of fundamental frequency (p<0.05, Wilcoxon sign rank test; mean across birds of +20%; range -19 to +43%). These data suggest that it is unlikely that changes to syllable structure might have arisen due to accidental damage to lMAN. (B) Percent change in mean fundamental frequency after mMAN lesions vs mean fundamental frequency before mMAN lesions.

**Figure 2 – figure supplement 4:**
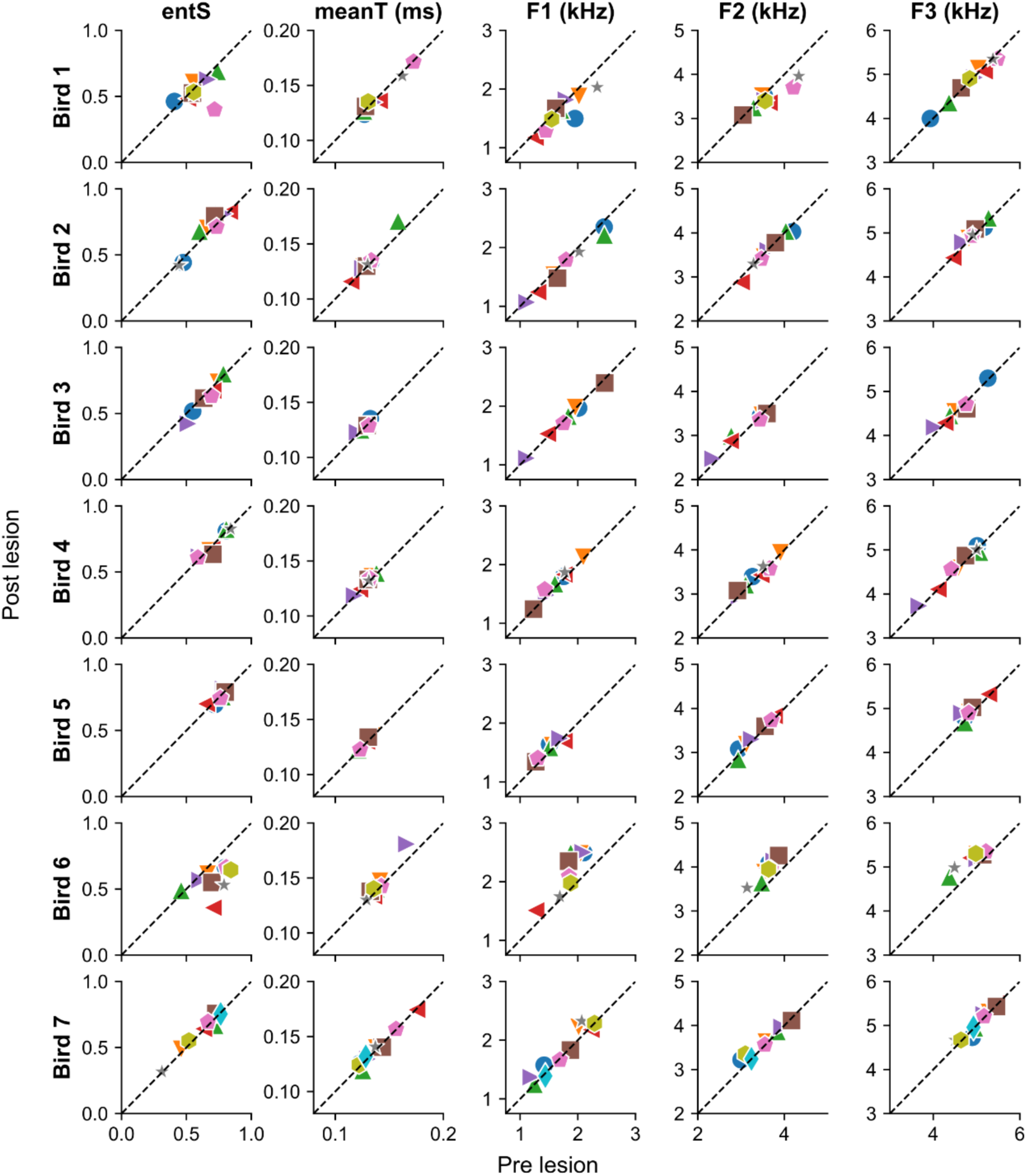
Selected acoustic features for all syllables in all birds before and after mMAN lesions. Different colors represent different syllable types per bird. ‘entS’ = Entropy of spectral envelope, ‘meanT’ = Temporal centroid for temporal envelope, ‘F1’ = First formant, ‘F2’= Second formant, ‘F3’ = Third formant. Acoustic features generally showed little change between pre and post lesion songs. They highlight as relative outliers the same individual examples that stand out in the average spectrograms in Figure 2 – figure supplement 1.

**Figure 2- figure supplement 5:**
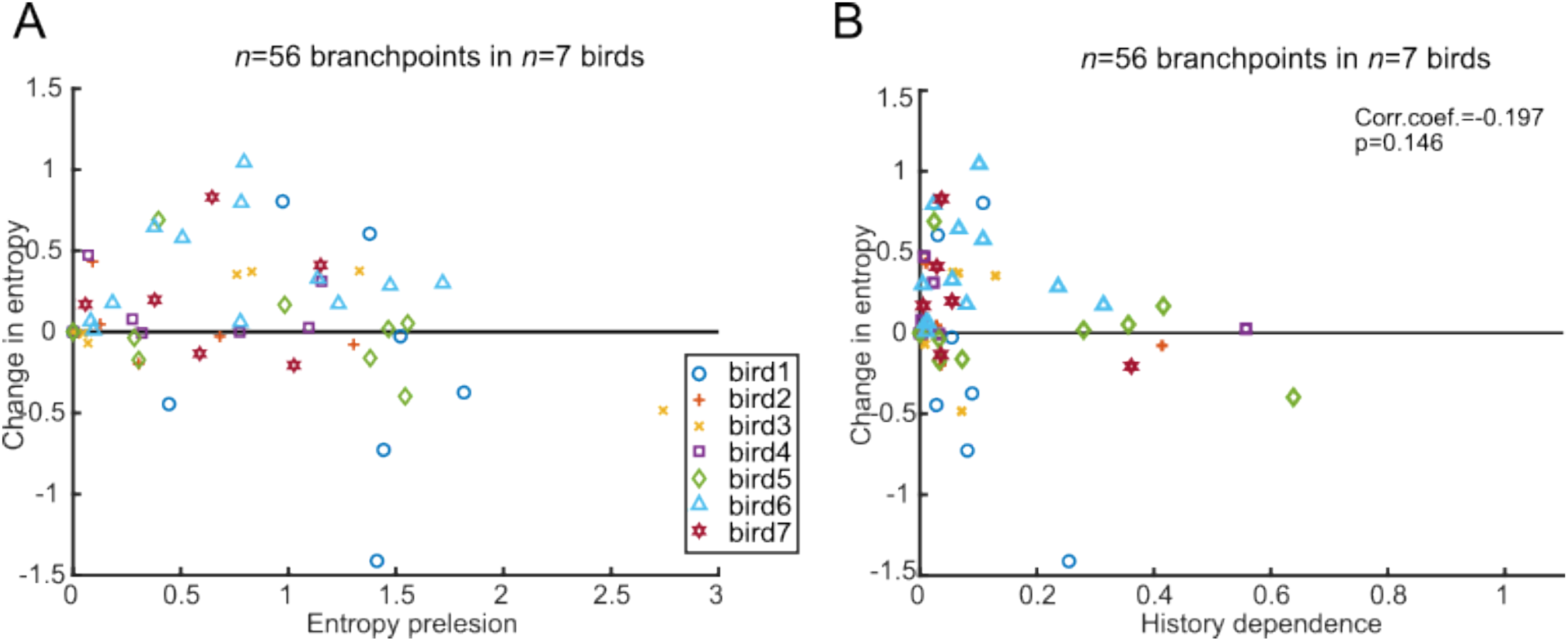
Change in transition entropy for branchpoints (A) Transition entropy for all individual branchpoints after mMAN lesions. (B) Branchpoints with low history dependence before mMAN lesions showed a nonsignificant trend towards higher changes in transition entropy.

**Figure 3 – figure supplement 1:**
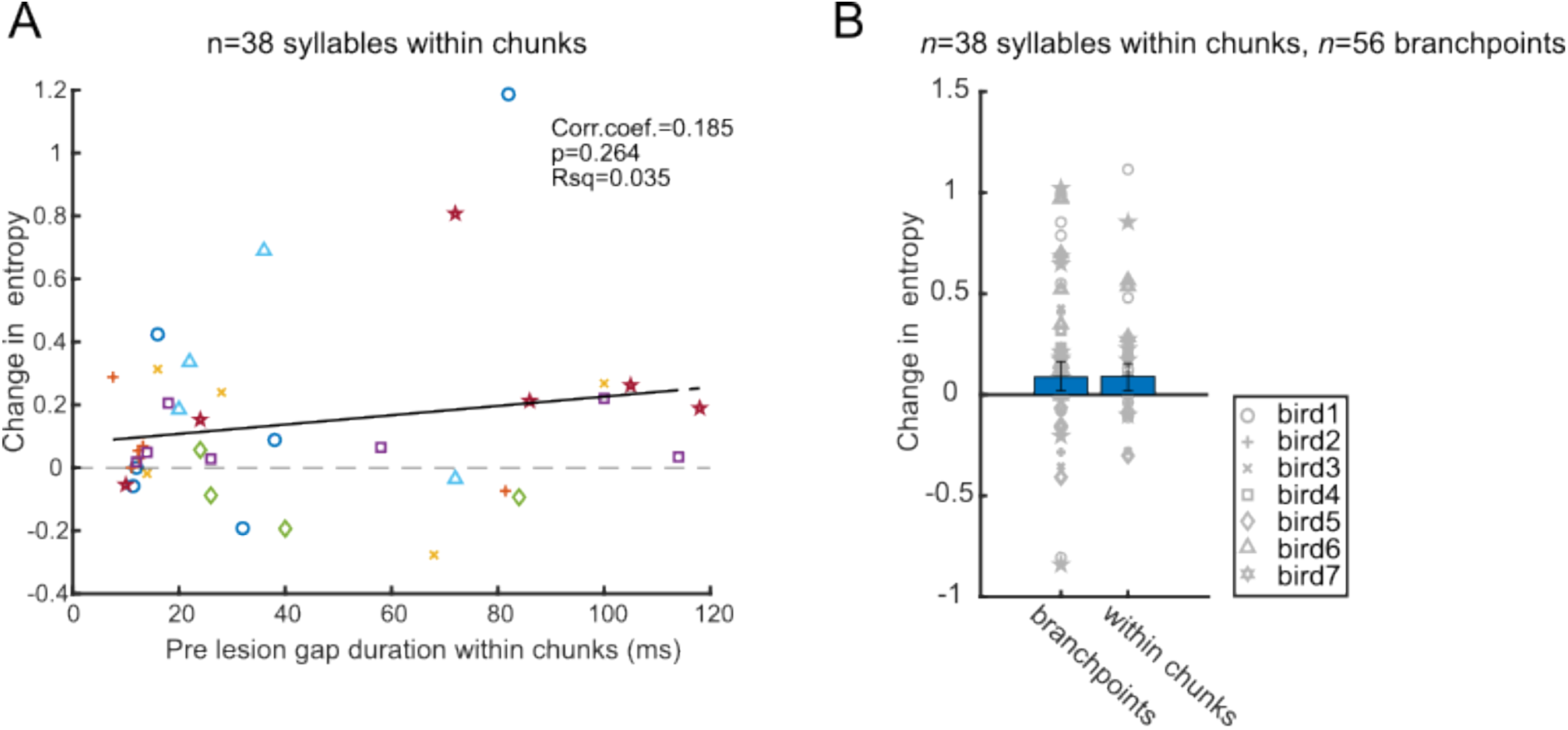
Change in transition entropy for transitions within chunks vs. branch points (A) Duration of gaps within chunks before mMAN lesions was not significantly correlated with entropy change at the corresponding transition (p> 0.05, Wilcoxon rank sum test). (B) Change in transition entropy was not significantly different for transitions within chunks and at branchpoints (p> 0.05, Wilcoxon rank sum test).

